# The heat-ramp method to study regulated cell death in a pathogenic yeast *Cryptococcus neoformans*

**DOI:** 10.64898/2026.01.19.700367

**Authors:** Madhura Kulkarni, Yining Liu, Quanxi Cheng, Chengzhang Zhu, Kimberly Kuhn, Ao Shen, Bingnan Jin, Arturo Casadevall, Heather M. Lamb, Zachary D. Stolp, J. Marie Hardwick

## Abstract

Human fungal pathogens cause a significant public health burden. While no reliable surveilence data are available, estimations suggest that 1 billion infections and over 2 million deaths are attributable to fungal infections annually worldwide. This drove the World Health Organization to generate a priority list of fungal pathogens for reearch, which includes the yeast *Cryptococcus neoformans* in a top critical priority. With the rise of drug-resistance and emerging fungal pathogens, new conceptual strategies for antifungal therapies are needed in addition to existing antibiotic development pipelines to meet clinical needs. Intrinsic cell death pathways encoded by pathogenic fungi are largely unstudied but could be leveraged for antifungal therapy analogous to anti-cancer therapeutics that activate apoptosis or other cell death mechanisms. Thus far, molecularly defined fungal cell death mechanisms are best characterized for only a few, predominantly model filamentous species. To extend these studies to pathogenic yeast, here we describe and demonstrate a tunable heat-ramp stimulus that when applied to small volumes of yeast cell suspensions reveals a protracted cell death process in the pathogenic yeast *Cryptococcus neoformans*. This low cost protocol induces robust and reproducible phenotypes to study gene-dependent mechanisms in laboratory strains and clinical isolates.

## 1. Introduction

The identification of self-execution mechanisms in microbes that resemble genetically defined cell death mechanisms in mammals emphasizes the conservation of cell death processes across species (1). Self-orchestrated cell death is facilitated by factors produced by the dying cell. This type of cell death is distinct from unregulated necrosis or loss of integrity due to direct assault, such as rapid cell breakage (e.g. lysate preparation) that does not involve genetically programmed mechanisms. The field of microbial cell death has entered a new era with the identification of multiple bacterial and several fungal executioners such as the gasdermin-like proteins that resemble mammalian innate immune and pyroptosis mediators (2-4). Self-death via membrane pores formed by bacterial and filamentous fungal gasdermins can protect their greater communities from the spread of phage infections or other predatory nucleic acids (5,6), and dying fungal cells are thought to be a crucial source of nutrients for survivors (7,8). Allogeneic rejection leading to fungal cell death is best characterized in non-pathogenic model filamentous fungi and is mediated by diverse molecular mechanisms, only some of which resemble mammalian mechanisms (6,9). However, semblances of known cell death executioner proteins (e.g. gasdermins) have not been identified in yeast species, suggesting that additional novel mechanisms are responsible.

Therefore, it is important to develop assays capable of identifying genetically regulated cell death mechanisms in pathogenic yeast of significant public health importance (10-12). We developed several assays designed to detect gene-dependent cell death in baker’s/brewer’s yeast *Saccharomyces cerevisiae*. The most powerful of these assays is the heat-ramp cell death assay (13-15). The key feature of this assay is that the heat-ramp stimulus is sufficiently mild to avoid death by direct assault (heat-shock) and yet cause death of almost all cells to prevent survival via normal stress responses. The heat-ramp assay was successfully applied to genome-wide screens in yeast and bacteria, identifying candidate pro-death factors and potential virulence factors (16-18). Genetic susceptibility or resistance to heat-ramp often reflects results obtained with alternative death-inducing stressors such as acetic acid, H_2_O_2_, and yeast killer viruses (13-15,19).

Here we describe the heat-ramp methodology adapted and updated for the opportunistic pathogen *Cryptococcus neoformans* (basidiomycete) (**Fig 1A**). A key aspect of the heat-ramp cell death assay is tunability to accommodate multiple growth conditions and species, robustly distinguishing the relative sensitivity of *C. neoformans* versus *Saccharomyces cerevisiae* (**Fig. 1B-E**). The goal of fine-tuning the death stimulus is to avoid gene-independent unregulatable death that occurs soon after treatment while simultaneously avoiding sublethal doses of stress that can be rescued by upregulation of stress responses, such as ESCRT-mediated repair of gasdermin pores in mammalian cells (20). Using the heat-ramp methodology, a genome-wide screen of ∼5000 knockout strains of *S. cerevisiae* (ascomycete) uncovered highly death-resistant strains lacking a potential death effector gene. This screen identified all four subunits of the yeast AP-3 vesicle trafficking complex. Further evidence indicates that AP-3 trafficks one or more cargo proteins to the vacuole that subsequently promotes cell death potentially through vacuole membrane permeabilization (18). This death-promoting function of AP-3 appears to be conserved in human pathogen *C. neoformans* (18), representing a novel role for AP-3 in regulating cell death not previously described in other organisms. However, vacuole and autophagy-related functions have critical roles in fungal cell death of plant pathogens including *Ustilago maydis* (basidiomycete) and *Magnaporthe oryzae* (ascomycete), which are of significant agricultural importance (21,22).

**Figure 1.**
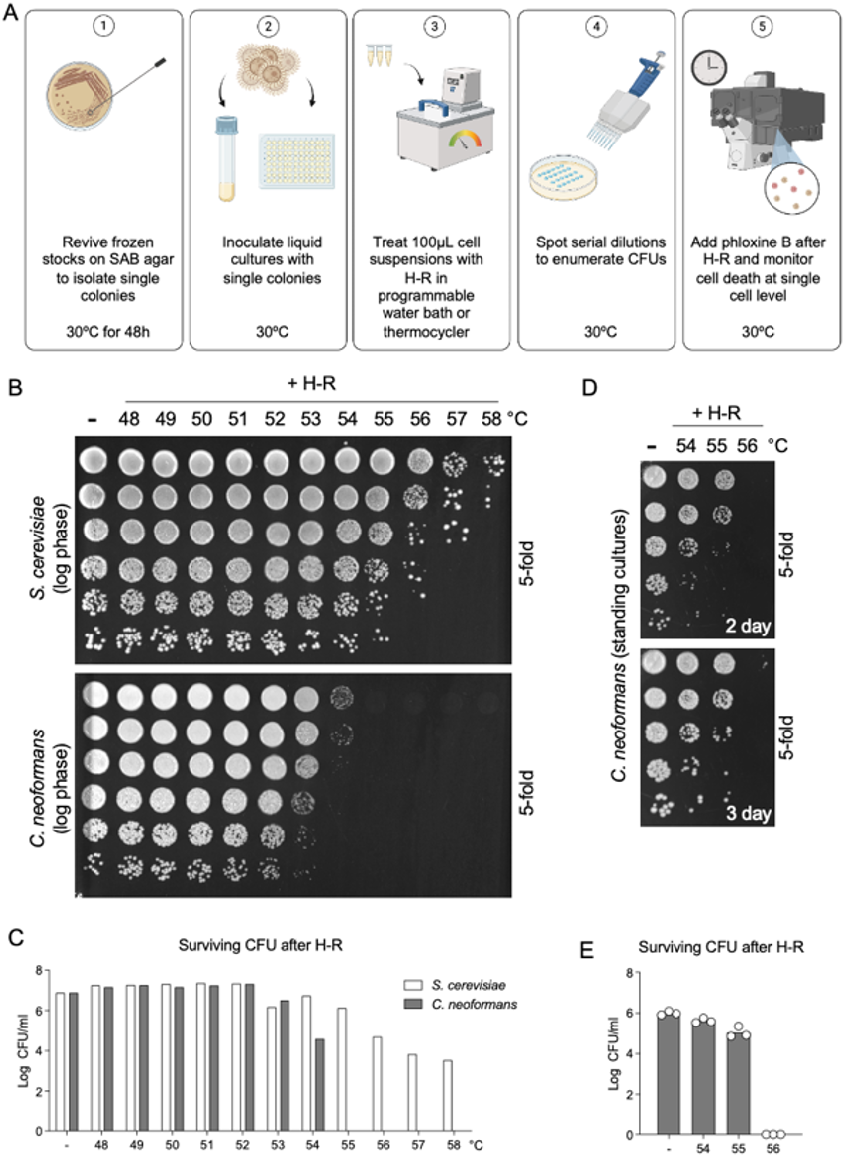
Application of heat-ramp stress using a programmable water bath induces cell death and loss of colony forming units (CFUs). (**A**) Graphical outline of the heat-ramp (H-R). The H-R assay is a treatment of gradually increased temperature (programmed in 1◦C increments), tunable to accommodate diverse strains of yeast as well as strain backgrounds, culturing methods and more (images generated in Biorender.com) (**B**) Dose-dependent H-R results based on growth of yeast, plating undiluted in first row followed by 5-fold serial dilutions (clonogenic survival) after a standard linear heat-ramp from 30°C to indicated maximums over ∼10 min for rolled log phase cultures of *S. cerevisiae* (BY4741) and *C. neoformans* (KN99α), revealing their known differences in temperature sensitivity. **(C)** CFUs quantified for panel B. (**D**) Standing cultures in 96-well format have higher viabilities and allow for ease of screening many strains. Quantification of CFU/ml at ≤ 2 days helps avoid merged colonies by day 3 of plate incubation (30°C) following H-R treatment. Undiluted samples in C and D were spotted in the first row followed by 5-fold serial dilutions (clonogenic survival). Based on prolonged incubations of spotted plates, undetectable colonies at 48 h may appear by 72 h or later but no significance differences were detected as these events are rare (not illustrated here). (**E**) CFUs quantified for 3 independent experiments shown in panel D.

The heat-ramp assay is not a predictor of thermotolerance (the ability to grow at warmer temperatures), and is not a heat shock assay that induces sudden death by assault. Instead, the heat-ramp methodology induces a protracted fungal cell death over many hours, consistent with an ongoing dying process. This was previously demonstrated for *S. cerevisiae* by detecting dying/dead cells with phloxine B staining, which is capable of detecting onset of cell death before other reagents we have tested, ensuring that the heat-ramp stimulus does not result in rapid (unregulated) cell death (18). Similarly, by plating phloxine-stained cells on agar and imaging at several time points after heat-ramp (**Fig. 2A**) or by automated video microscopy (**Fig. 2B**), we observe a slower time-to-death for *C. neoformans* (24-36 h depending on conditions) compared to *S. cerevisiae* (∼16 h) (18). Thus, heat-ramp assay parameters can be easily adjusted to accommodate different metabolic states, genetic backgrounds and levels of sensitivity (14,15,18). This method applied to *C. neoformans* identified cell death resistance in mouse-passaged brain isolates compared to parental strains (23). The heat-ramp assay also distinguished an emerging fungal pathogen *Rhodotorula mucilaginosa* collected from an urban heat island compared to the same species collected from a cooler urban location a short distance away (24). Resistance to cell death in the context of infection may permit increased replication and dissemination of disease. Therefore, assays to detect death-resistance and to map resistance mechanisms can support efforts to develop antifungal treatments with clinical focus.

**Figure 2.**
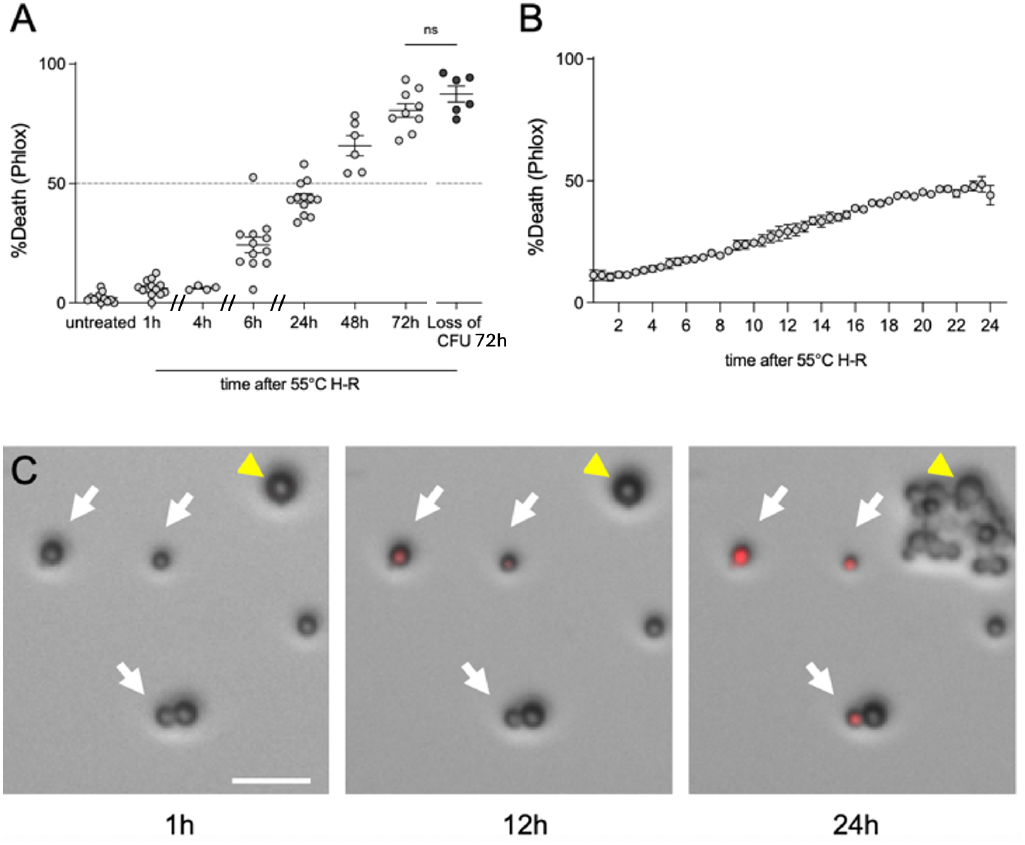
Slow death kinetics of *Cryptococcus neoformans* after heat-ramp detected by fluores-cent vital dye at early time points. (**A**) Compilation of independent heat-ramp assays (30-55°C ∼10 min) z video microscopy for single cell analysis of phloxine-stained KN99α compiled and compared to loss of CFU at 72 h after treatment. Death quantified as % of starting viable cell number stained with phloxine (not counting progeny of surviving cells as shown at 24 h). Time-to-death 50% was ∼38 h, (linear regression). No significant difference found comparing incubation of phloxine-stained cells to unstrained CFUs (counting visible colonies, p=0.145 (two-tailed T test). Loss-of-CFU calculated as [(colony# post treatment)/ (colony# of untreated control) x100]. (**B**) Automated analysis of phloxine uptake during live cell imaging, analyzed by ImageJ. Mean ±SEM per data point from 2 independent experiments (Leica widefield microscope). (**C**) Frames from video show phloxine uptake (death) at delayed times after heat-ramp. Cells that will die in 24 h (white arrows), a single survivor proliferates by 24 h (yellow arrow). Scale bar 20 µm

This assay is relatively simple, reproducible, and low-cost, making it accessible to a wide range of laboratories without the need for highly specialized equipment. In its simplest form, the heat-ramp assay can be successfully achieved with only a standard water bath, visible spectrophotometer, micropipette, and basic microbiological supplies. Reproducibility ensures that results can be directly compared across different strains, clinical isolates, independent experiments, and laboratories.

Before embarking on the heat-ramp assay, there are several key requirements for success. Foremost, it is essential to use a heating machine that is capable of delivering heat evenly to all samples. For example, all models of thermocycles sold in 2023 that we tested exhibited pronounced non-uniform well-to-well heating, each machine displaying a distinct and reproducible heating pattern that was far beyond what could be compensated by batch corrections (**Fig. 3**) (25), unlike our older Mastercycler machine with much smaller variations amenable to batch correction by well (18,26).

**Figure 3.**
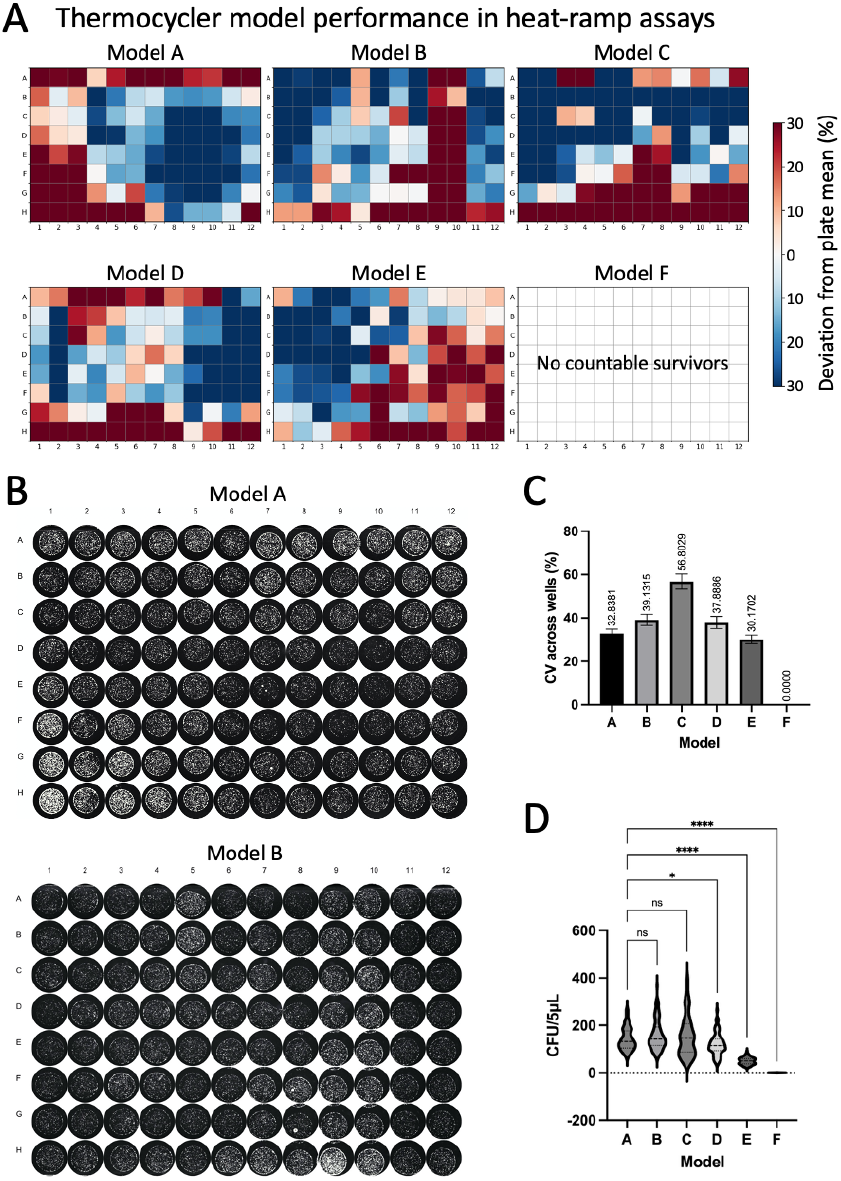
Example thermocycler heat-blocks exhibit reproducible signatures across 96 wells revealed by the heat-ramp assay. (**A**) Heatmaps showing a wide range of well-to-well variation in yeast survival due to highly reproducible but uneven heating patterns in 8 different thermocyclers, 6 different models from 4 vendors: Eppendorf model-A (1 of 3 Mastercycler® X50 tested, with individual well counts ranging from ∼30-300 microscopic colonies per spot; three models-B,C,D from Applied Biosystems: Veriti, VeritiPro, and Proflex, respectively; model-E: Biometra TAdvanced from Analytik Jena; model-F: BioRad C1000 Touch. The same solution of *Saccharomyces cerevisiae* was seeded into all 96-wells, treated with the same heat-ramp program, plated to determine cfu in multiple experiments. Color scale represents deviation from the plate mean CFU. Blue: lower survival (lower heat); red: higher survival (higher heat). (**B**) Example heat-ramp assays imaged and enumerated on a CTL Biospot Reader. Each machine had a reproducible distinctive signature pattern of heating that readily identified individual machines of the same model # (not shown). (**C**) Coefficient of variation (CV) of CFU counts in 96 wells reveal high well-to-well variance. Model F killed all yeast, CV = 0. (**D**) Violin plots of CFUs illustrate inter-plate variability. Data represent mean ± SD (N = 2-9 independent experiments per model). One-way ANOVA based on plate-averaged CV values; ns, not significant; p < 0.05 (*); p < 0.0001***.

When interpreting results, a key point to recognize is that the sensitivity of an optimized heat-ramp assay facilitates detection of variant phenotypes within a population. Seemingly inconsistent results between independent experiments are often misinterpreted as technical issues, but instead are genuine variant phenotypes due to mixed genotypes within a population, as demonstrated for *Saccharomyces* (16). This problem is resolved by testing side-by-side several different colonies picked from a streaked plate. Alternatively, for sticky strains of *Cryptococcus*, it may be necessary to microdissect single cells to effectively separate genotypes within a population.

## 2. MATERIALS

### 2.1 Critical equipment and certification to acquire ahead

1. Machine capable of precisely heating multiple 50-100 μl samples evenly (see *Notes 1-3*)
  a. Water bath with programmable temperature control, strong water circulation and reliable thermometer (see *Notes 4-8*)
  b. Thermocycler – validated to heat evenly with narrow temperature variance and programmed to linearly ramp (programmed in 1◦C increments) to desired temperature over 15-20 min as determined to be optimal (**Fig. 3**) (see *Notes 9 - 10*)
2. Spectrophotometer (e.g. Thermo Scientific Genesys 10S Vis) or plate reader
3. 30◦C incubator with a humidity box or tray (*see Note 11*)
4. Roller drum (TC-7, New Brunswick Scientific) with tube rack positioned at ∼45° angle for incubating cultures tubes for log phase assays
5. Fluorescence microscope equipped with incubator and humidifier for video microscopy time course (e.g., Leica widefield microscope)
6. Calibrated multichannel pipettes and matching tips
7. Biosafety Level 2 laboratory approved for *Cryptococcus neoformans*

### 2.2 Yeast and Culturing Supplies and Reagents for Heat-Ramp Assay (*see Note 12-13*)

1. Yeast strains, such as KN99α, H99 (e.g. FGSC https://www.fgsc.net/ or ATCC) or low passage clinical isolates from deidentified clinical banks]
2. Sabouraud (SAB) liquid broth (BD Difco), autoclave sterilized (*see Note 14*)
3. YPD liquid medium: 2% peptone (Peptone-Y, MP Biomedicals), 1% yeast extract (Fisher Scientific), and 2 % glucose (J.T. Baker), autoclave sterilized.
4. Round bottom polypropylene culture tubes with loose-fitting caps (e.g., 12 ml, Falcon) for log phase assays
5. 96 well micro plates, flat bottom, non-treated for culturing multiple strains
6. 0.2 ml PCR tubes

### 2.3 Colony Forming Assays

1. 96-well microplates (flat-bottom, 200 μl)
2. Omnitray plates 128 × 86 mm (or 10 cm dishes can suffice) (ThermoFisher)
3. 10 cm dishes
4. SAB agar for plates/dishes [3% liquid SAB broth, 2% agar (BD)], autoclave sterilized
5. Sterile inoculation loops

### 2.4 Tracking cell death by fluorescence microscopy

1. Phloxine B (Fisher, final concentration 2 μg/ml) (*see Fig 2*)
2. SAB agar poured thinly (∼15 ml) in Omnitrays to facilitate inverted microscopy imaging

## 3. METHODS

### 3.1 Heat-Ramp Cell Death Assay (see *Note 15 and 16*)

1. Yeast cells from frozen −80°C glycerol stocks (without thawing) onto SAB 10 cm agar plates and incubate at 30°C for 48 hours. (*see Notes 17 and 18*)
2. Transfer perfectly symmetrical single colonies of similar size from the plate with a sterile loop or other implement into 200 µL SAB (or YPD) in flat bottom 96-well plates for overnight incubation of standing cultures (∼16-22 h) at 30°C. (*see Note 19-21*)
3. Alternatively, disperse single colonies in 2 ml liquid medium in sterile culture tubes and incubate overnight (18-24 h) at 30°C on a roller drum positioned at an upward angle and rotation rate ∼35 revolutions/min. (*see Note 14 and 18*)
4. Before counting, cell clumps must be dispersed vigorously by trituration. (*see Note 14 and 18*)
5. Measure cell concentration immediately of a 1:10 dilution at OD_600_ in a cuvette using a spectrophotometer (plate reader or counted by hemocytometer).
6. Adjust all cultures to OD_600_ 0.1 (∼3x10^6^ cells/ml) in 200 μl fresh SAB liquid medium and incubated at 30°C without shaking.
7. Before performing the heat-ramp assay, bring agar plates to room temperature and ensure they are dry to avoid merged spots of serial dilutions. (*see Note 22*)
8. At every step, cultures need to be gently resuspended before dispensing or diluting as cells settle quickly, which can cause major inconsistencies in the results. (see *Note 23*)
9. Normalize each culture to OD_600_ = 0.5 in fresh medium with sufficient volume for 100 µL per treatment group (untreated and heat-ramp treated). (see *Note 23*)
10. Transfer 100 μl of each culture to 0.2 ml PCR tubes using a calibrated multi-channel pipet and place in a programmable water bath with consistent water level (volume) and circulation equilibrated to 30°C and reset temperature to 54°C or 55°C; adjust as needed depending on whether water bath heating slows as it nears maximum temperature. (see *Note 23*)
11. Alternatively, use a validated programmable thermocycler to apply a heat-ramp stress (see Fig. 3). Thermocycler programs should hold at 30°C for 45 seconds to equilibrate, and ramp from 30°C to final maximum temperature (e.g., 54°C or 55°C) over approximately 10 minutes (the exact time will vary depending on the exact volume of water). (see *Note 23*)
12. The remaining untreated OD-adjusted cultures are serially diluted (5-fold) in sterile deionized water using a multi-channel pipet and plated on SAB agar plates as controls to verify equal cell numbers prior to treatment.
13. Treated cultures are mixed and serially diluted (5-fold) and 5 µl spots are plated on SAB agar plates to compare surviving CFUs to untreated samples. Keep consistent timing between heat-ramp in medium and dilution in water (e.g. 15 - 75 min). (see *Note 15*)
14. Spots are dried with plates lidded and then inverted and incubated at 30°C. (see *Note 22*)
15. Image plates under magnification and careful focusing to enumerate CFUs 1.5 to <2 days. If not using automated colony counting (e.g. CTL biospot reader) (18), rather than counting CFUs by eye (which varies person-to-person) (14), counting well-focused plate images enlarged on screen at matching magnifications is recommended to distinguish merging colonies, and avoid counts less than 8-10 colonies per spot for quantification. (*see Note 24-31*)
16. Alternatively, determine cell viability at specified early times after heat-ramp (before cell proliferation of survivors) by imaging phloxine-stained aliquots on thin SAB agar plates, counting microscopy images, ∼300 cells per sample, per time point, taking care to collect data from representative fields. (*see Note 32*)

### 3.2 Tracking cell death at a single-cell level following heat-ramp

1. Stain heat-ramp treated cells in 2 μg/ml of phloxine B dye and spot dilutions immediately on thin SAB agar plates that are compatible with fluorescence imaging (e.g. OmniTray plates). (see *Note 32*)
2. Monitor fluorescence and bright field images by video microscopy every 10-30 minutes over 12-30 h. Using a Cytation imager or video microscopy ensures that the progeny of survivors after heat-ramp are not counted as starting cell numbers (see Fig. 2C, right panel).

## 4. Notes

1. Critical pretesting of heating machines is required to ensure even heating of cell samples, achieved by placing equal aliquots of the same homogeneous culture into each well/tube and performing a heat-ramp assay (see **Fig. 3**).
2. Additional pretesting to calibrate the heat-ramp assay conditions by testing timing at each step, maximum temperature and potentially variable ramp rates to maximize reproducibility based on available equipment (14).
3. To maximally reveal heat-ramp phenotypes in different yeast strains may require gentler (e.g. weaker phenotypes) or stronger treatment conditions (e.g. strongly death-resistant strains) (see **Fig. 1B**).
4. For water bath heat-ramps, ensure that the volume of water in the water bath is adjusted to the same level for every experiment to improve reproducibility, e.g. use a level to score the outside of a clear plastic water bath to mark preferred water level.
5. Preheat the water bath to 30°C before retrieving overnight cultures from incubator to work as efficiently as possible to minimize lag times between steps.
6. Ensure that the cell samples in floating tubes in a water bath are fully and uniformly submerged and at a maximally achievable distance below the water surface. A floating tip rack is useful to submerge the bottom of tubes without submerging the lid.
7. Avoid lag time in removing cultures from the water bath by using strip tubes or place tape across tube tops to remove all tubes from the bath simultaneously.
8. After retrieving tubes from the water bath gently dry off PCR tubes with absorbent tissues to ensure that no water is around the caps before opening the lids.
9. For thermocycler heat-ramps, perform initial assays to determine the consistency of heating across the heat block at different maximum temperatures by placing aliquots of a single yeast culture in all 96 wells (see **Fig. 3**).
10. For smaller experiments using the thermocycler, select specific positions in the thermocycler heat block where repeated pretesting results indicate that similar results are achievable. Recheck these positions periodically to ensure reproducibility.
11. Recommend keeping overnight 30°C incubations of plates and multi-well cultures in a humid environment to avoid evaporation and edge effects.
12. Equivalent consumables and equipment are available from a variety of alternative providers.
13. Recommend using the same lot numbers of media components for large screens.
14. SAB medium is recommended and may differ from phenotypes in YPD. Rolled overnight culture tubes in SAB, unlike YPD, typically result in ∼50% inviable cells, in contrast to 96-well standing SAB cultures with low single digit % death).
15. Technical consistency between experiments can be improved by controlling multiple steps, including but not limited to expert multi-channel pipetting, consistent times between steps, matched ODs (or use hemocytometer for unclumped cells) at start and end of overnight incubation to match starting cell number and other factors (as longer growth phases can lead to increased or decreased death resistance depending on cell density and rolled or standing incubations), accuracy of OD adjustments depends on disruption of cell clumps, and timing between the multiple steps when setting up the heat-ramp assay and the time between heat-ramp and plating. Allowing cells to recover for specified time (e.g. 30 min or 1 h) in medium before preparing serial dilutions in water will increase survival (e.g. heat-ramp in water instead of medium greatly reduces cell viability).
16. Triplicate overnight cultures for each sample can reduce variation due to pipetting errors in CFU plating per experiment.
17. Recommend assuming that all passaged stocks, especially of clinical isolates and deletion strains are composed of heterogeneous phenotypes, as gene-deletion-driven genome evolution has been demonstrated for *Saccharomyces cerevisiae* and likely similar for *Cryptococcus neoformans* (16). For example, deletion of genes for different components of the same protein complex often evolve mutations in the same secondary gene or complementation group (16). Therefore, it is critical to analyze single-cell-derived colonies, which is harder to achieve for clinical isolates of *Cryptococcus neoformans*.
18. Thorough mixing when diluting yeast cultures and when disrupting cell clumps is critically important, but it is also important to avoid generating air bubbles. Recommend checking successful clump disruption by examining a small drop of cells under a light microscope (especially before using a hemocytometer to count yeast cells).
19. Streaking plates with only a few swipes is unlikely to produce single cell-derived clones of *Cryptococcus neoformans*. However, success is more likely if plates are streaked thoroughly, including backtracking over the same streaks in attempt to separate cells.
20. However, if variable phenotypes persist, this is likely because the population remains heterogenous (16). In this case it will be necessary to microdissect individual cells using a tetrad dissection scope with microfiber needle (and some practice).
21. For microdissection, on Day 0: divide/mark a SAB agar plate into two areas, two-thirds of the plate for a holding area and one-third for a picking area. Streak *Cryptococcus neoformans* from frozen stocks in the picking area, and incubate at 30℃ overnight (to allow recovery and begin regrowing). On Day2: use a tetrad needle with a flat end to tease apart a cluster of yeast cells under microscope and array the single cells in the holding area. Verify that each contains only a single cell; mark exceptions. Incubate at 30°C for 48 h.
22. Agar plates need to be dry before storing and may need further drying before use. Ensure agar plates are dry (no condensation visible and typically left to solidify on level benchtop for ∼2 days before storing in 4°C) before spotting dilutions of cultures to ensure spots dry and do not touch.
23. After OD normalization, the heat-ramp treatment should be started immediately as delay allows refed cells to start growing, and result in different responses to the heat-ramp treatment.
24. Counting visible CFUs early (∼36-42 h post treatment) helps avoid overcrowded colonies
25. Employ consistent rules when counting colonies in spot dilution series. Often colonies can merge and colonies that are not round, or show indentations between multiple peaks should be counted as separate colonies. Or use a biospot reader capable of counting microscopic colonies (18).
26. Consider stickiness of different strains to avoid undercounting.
27. To plate in straight lines with a multichannel pipet, use a 96-well pipetting template underneath agar plates to help with spotting dilutions (e.g. empty 96 well tip rack or dots printed on paper in 96 well format).
28. Pipette tips should be firmly pressed into the multichannel pipette to ensure accurate volume delivery.
29. Depress multichannel pipette to have cultures suspended at the tips and then gently touch to agar plate to spot the 5 ul, as opposed to risking the pipette tips touching the agar and risk bubbles when dispensing/spotting.
30. Change tips at every step when preparing dilutions, mixing well at each step. When plating/spotting the serial dilutions, start from the most diluted (bottom row) and work backwards (up) to use the same tips. Mix well, cell settle quickly.
31. Beware of contamination; collect growth from yeast plates and examine under a microscope to detect low levels of bacterial contamination. Consider plating on ampicillin plates to clean up critical original stocks. Don’t open plate lids except briefly or perform plating experiments in a flow hood, but beware of excess plate drying.
32. Note that phloxine staining is more sensitive and can often detect dying cells several hours before propidium iodide staining is detected (18). At the recommended low concentration of phloxine, no measurable effects on viability were detected but this could potentially differ for other strains.

## 5. Concluding Remarks

This heat-ramp method uniformly delivers heat over several minutes to small volumes of yeast cells to induce protracted cell death of lab strains and clinical isolates of the opportunistic pathogen *C. neoformans*. Allowing cell death to occur over a prolonged time frame facilitates dissection of the molecular-cellular events that occur during the dying process. Yeast viability following heat-ramp treatment can be determined in real time on a solid surface at a single cell level using a vital dye, or determined as clonogenic survival (CFU). Both low and high throughput readouts (single tubes vs. 96-format) are relatively easy to perform, can be conducted using existing lab equipment, and allows for visualization of death in real time.

This powerful assay can reveal hidden heterogeneity otherwise challenging to detect. The heat-ramp assay can be used to evaluate variation in cell death resistance within a population and can be used iteratively to isolate phenotypes otherwise masked within a yeast population. This standardized assay allows quantification and direct comparison between different strains. Microdissection of single cells reduces the confounding influence of mixed subpopulations and allows resolution of inter-colony variation that might otherwise obscure genetically determined cell death phenotypes. The combination of CFU readout and time-lapse imaging allows discrimination between immediate (unregulated) and delayed cell death (evidence of an ongoing process).

## Acknowledgements

We would like to acknowledge members of the Casadevall lab for their assistance and guidance.

